# VETA: a Matlab toolbox for the collection and analysis of electromyography combined with transcranial magnetic stimulation

**DOI:** 10.1101/610386

**Authors:** Nicko Jackson, Ian Greenhouse

**Author notes:** **Corresponding author:** Ian Greenhouse, Address: Department of Human Physiology, University of Oregon, 181 Esslinger Hall 1525, University St. Eugene, OR 97403.

## Abstract

The combination of electromyography (EMG) and transcranial magnetic stimulation (TMS) offers a powerful non-invasive approach for investigating corticospinal excitability in both humans and animals. Acquiring and analyzing the data produced with this combination of tools requires overcoming multiple technical hurdles. Due in part to these technical hurdles, the field lacks standard routines for EMG data collection and analysis. This poses a problem for study replication and direct comparisons. Although software toolboxes already exist that perform either online EMG data visualization or offline analysis, there currently are no openly available toolboxes that flexibly perform both and also interface directly with peripheral EMG and TMS equipment. Here, we introduce Visualize EMG TMS Analyze (VETA), a MATLAB-based toolbox that supports simultaneous EMG data collection and visualization as well as automated offline processing and is specially tailored for use with motor TMS. The VETA toolbox enables the simultaneous recording of EMG, timed administration of TMS, and presentation of behavioral stimuli from a single computer. These tools also provide a streamlined analysis pipeline with interactive data visualization. Finally, VETA offers a standard EMG data format to facilitate data sharing and open science.

## Introduction

Ever since the groundbreaking studies conducted by Luigi Galvani in the late 18^th^ century, the role of electricity in the control of muscles has been an important and fruitful topic of research. Electromyography (EMG) is a powerful method for recording muscle electrical activity that yields rich data and has a wide variety of applications. In combination with other techniques, EMG provides unique insights into the fundamental mechanisms that drive animal and human movement.

Transcranial magnetic stimulation (TMS) is a form of noninvasive brain stimulation that when combined with EMG can measure corticospinal excitability with high temporal resolution. A single TMS pulse of sufficient intensity administered over a muscle representation in primary motor cortex can excite the corticospinal pathway and activate the targeted muscle to produce a motor evoked potential (MEP), which reflects the momentary state of excitability of the output pathway of the motor system. In addition to commonly evaluated features of EMG activity associated with voluntary muscle contraction, many other metrics can be quantified using the combination of EMG and TMS such as MEP onset latency, MEP peak-to-peak amplitude, and MEP duration (for review see Bestmann and Krakauer 2015). These metrics are often used to characterize the dynamics of motor system excitability at rest and during the performance of behavioral tasks.

TMS derived metrics have a wide range of research and clinical applications in the fields of Physiology, Psychology, Psychiatry, Kinesiology, Biomechanics, Neuroscience, and Neurology. The number of research studies utilizing combined EMG and TMS methods has increased dramatically in recent years. According to PubMed, in the 20 years preceding the writing of this article there have been 4955 published studies that include both the terms “electromyography” and “motor evoked potential” anywhere in their text. This includes a wide variety of research studies with neurological and neuropsychological applications ranging from the evaluation of basic mechanisms, e.g. assessing the speed of nerve conduction and excitability state of the motor system in Parkinson’s disease (Kandler et al. 1990; Vacherot et al. 2010), to the prediction of clinical outcomes. For example, TMS elicited MEPs may help predict which patients are likely to recover after stroke (Smith and Stinear 2016) or are likely to respond to non-motor repetitive TMS for the treatment of depression (Oliveira-Maia et al. 2017). TMS has also shown promise for the diagnosis of Alzheimer’s disease because these patients exhibit heightened sensitivity to motor TMS (Di Lazzaro et al. 2004). In non-clinical research, the combination of EMG and TMS has been applied to the assessment of biomechanical performance, e.g. the study of relationships between limb position and motor system excitability (Lin et al. 2015) and between cortical excitability and volitional control of gait (Ito et al. 2015). In the fields of Neuroscience and Psychology these methods have been applied to the investigation of a plethora of behavioral and psychophysical phenomena including changes in motor excitability during the preparation of movement (Duque et al. 2010; Greenhouse et al. 2015; Lebon et al. 2016), intrinsic individual differences in motor performance (Greenhouse et al. 2017), and volitional control of brain states (Ruddy et al. 2018). There are important veterinary research applications as well (Nollet et al. 2004).

While these wide-ranging research applications are not necessarily commonplace in the clinic, some clinical procedures for motor TMS are fairly common (Di Lazzaro et al. 1999; Hallett 2007). These include the determination of individual sensitivity thresholds for administering repetitive TMS therapies for the treatment of migraine (Barker and Shields 2017), major depression (O’Reardon et al. 2007), and other psychiatric disorders (Bersani et al. 2013), mapping the motor cortex to guide neurosurgery (Seynaeve et al. 2019), and evaluating the integrity of cortical and output pathways in motor disorders (Edwards et al. 2008). These procedures occur on a daily basis in clinics throughout the world.

Given the diverse applications of these methods, there is an apparent need for standardized and automated data collection and analysis tools. While some studies have relied on the visual detection of TMS elicited muscle movements, the determination of an individual’s sensitivity to TMS using this method alone overestimates their true threshold, and this has important safety considerations for both research and clinical applications (Westin et al. 2014). Standardization and automatization of combined EMG and TMS procedures can help to alleviate risks associated with over-stimulation by more accurately and consistently determining individual thresholds. Unfortunately, automated simultaneous EMG measurement and TMS administration has traditionally required interfacing multiple computers and the adaptation of software developed for purposes other than MEP measurement.

Open source toolboxes exist for separate online biosignal visualization (SigViewer, Brunner et al. 2013) and offline EMG analysis and visualization (CortExTool, Harquel et al. 2016; MAVIN, Mullins and Hanlon 2016). However, the application of separate software for online and offline procedures introduces extra data conversion steps which can produce errors or artefacts related to incompatibilities across platforms. Combining both online and offline functions within a single toolbox provides a number of advantages by streamlining the data collection and analysis process. Moreover, while there is a general consensus on guidelines for the safe and ethical administration of TMS (Rossi et al. 2009), there is no existing common standard format for EMG data recordings. The lack of a universal standard data format adds an additional challenge for comparisons across studies and creates an artificial divide between users of different software packages. This problem is compounded by a lack of tools for converting between the many different formats of EMG data. A standardized, automated method for recording EMG and calculating common metrics can facilitate robust comparisons across research studies and improve the overall reliability and safety of experiments. Additionally, tools for converting a variety of EMG data types to a common format will support open science practices.

Here, we describe the Visualize EMG TMS Analyze (VETA, https://github.com/greenhouselab/VETA) Matlab toolbox for on-line EMG data visualization, automated analysis, and interactive off-line data review. The VETA toolbox was specifically designed to interface with EMG and TMS equipment to facilitate the measurement of MEP and EMG activity patterns commonly observed in laboratory research studies and in the clinic. The code is written to be flexibly adaptable for new applications, includes tools for importing data from other software, and is freely available for download.

## Workflow

The VETA toolbox is comprised of three separate tools designed to be executed in sequence:

1. recordEMG
2. findEMG
3. visualizeEMG

First, recordEMG simultaneously visualizes and collects EMG data. Then, findEMG identifies common EMG events in the data and calculates metrics. Finally, visualizeEMG provides a graphical user interface for the interactive review of the raw data with event markers displayed alongside the calculated metrics. The following sections further describe the capabilities of each function and how to execute them.

**Figure 1.**
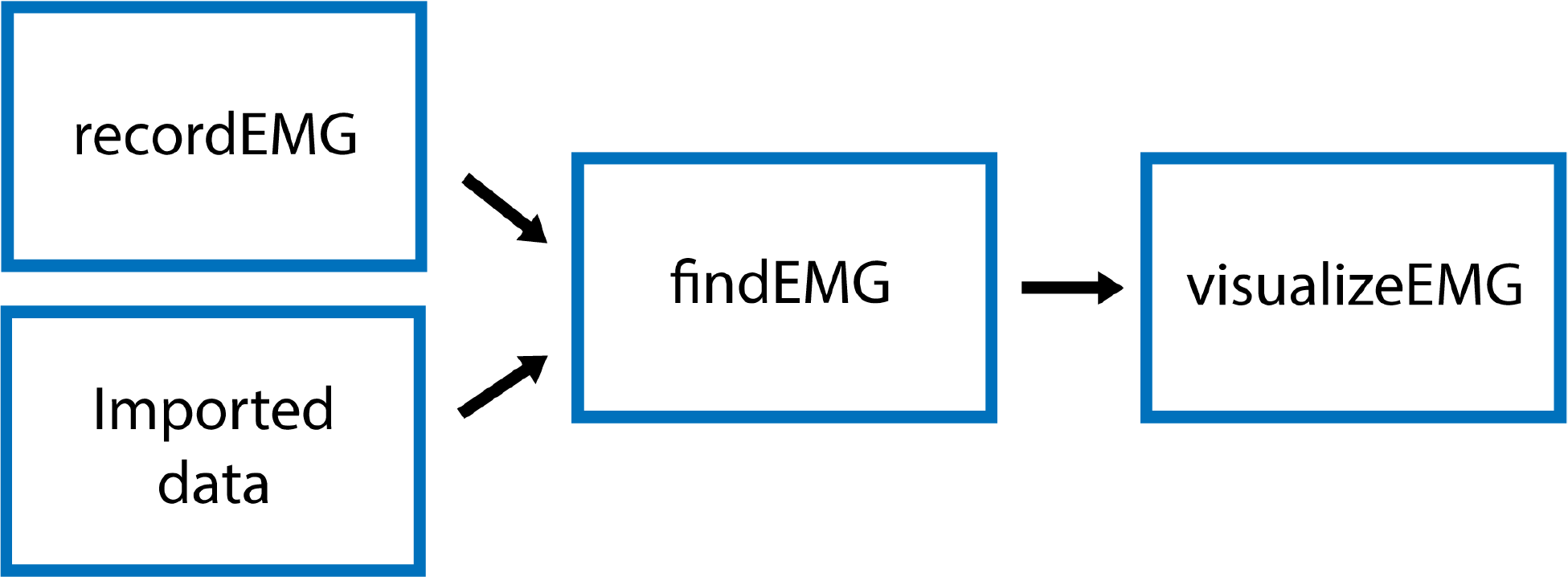
Schematic of VETA workflow. Data can either be collected with recordEMG or generated by other data acquisition software such as EMGworks, BIOPAC AcqKnowledge, or Signal. The automated detection of events is performed with findEMG and associated metrics are calculated. Interactive review of data is performed with visualizeEMG, allowing the user to manually correct misidentified events.

### 1) recordEMG

This function plots and records consecutive sweeps of EMG data.

#### Syntax

~~~
>>recordEMG
~~~

#### Description

Parameters (default values in parentheses) are specified below the header block and include the number of sweeps (n = 4000), sampling rate (5000 Hz), sweep duration (2 s), Y-axis range for plotting (±0.1 V), X-axis range for plotting (0 to 2 s), and reference lines for visualizing EMG event amplitudes (±0.025 V). All these values can be tailored to suit the needs of individual experiments. The default reference line values were selected based upon the standard approach used to determine MEP resting motor threshold, i.e. the lowest intensity TMS stimulus to produce MEPs of at least 50 μV on 50% of attempts.

There are two options for saving data using the ‘save_per_sweep’ parameter. Option 0 saves the data at the end of the complete acquisition session. With this option data are collected with minimal delay between sweeps. Alternatively, option 1 saves the data after each individual sweep. Saving the data after each sweep introduces an acquisition delay depending on the computer processor speed and RAM. The default ‘save_per_sweep’ setting is 0 resulting in minimal delay between consecutive sweeps.

The **recordEMG** function can also be used to calculate maximum voluntary contraction (MVC). Additional parameters are specified below the header block for the calculation of MVC. The default settings for measuring MVC include four sweeps at 4 s per sweep from a single channel. At the completion of recording, the maximum peak-to-peak EMG amplitude is calculated from the four consecutive sweeps. A calculated ‘MVC_percentage’ (5 %) relative to the maximum peak-to-peak EMG amplitude recorded across the four sweeps is outputted to the command line. This is particularly useful for setting up commonly used paired-pulse protocols, such as SICI (Kujirai et al. 1993; Ziemann et al. 1996), or cortical silent period (CSP, (Wilson et al. 1993; Werhahn et al. 1999) protocols during which muscle contractions are often maintained at a set percentage of MVC.

When **recordEMG** is called, the user is prompted to choose between practice (0) or run/save (1). When in practice mode, data is not saved. This mode is useful for visualizing data when storing the data is unnecessary, e.g. troubleshooting or evaluating EMG quality during initial setup for recording. In contrast, if run/save is selected, data is saved according to the method set by the ‘save_per_sweep’ parameter described above.

The user is then prompted to input whether maximum voluntary contraction (MVC) will be measured (1 for yes, or 0 for no). If MVC is measured, only one channel will be displayed, otherwise the user is prompted to enter the number of channels to record. Additional prompts request the user enter the subject ID, handedness, sex, date of birth, and resting motor threshold. These entries can be modified or commented out of the code according to the information the experimenter would like to store with the data. Altering these inputs or associated variables does not pose any loss of functionality to the EMG recording capabilities.

A ‘diode’ input parameter is also specified. Setting this value to 1 adds a supplemental recording channel to the data collection. This can be used to measure any additional input through a supported DAQ device that the experimenter would like to simultaneously acquire with the EMG. For example, a photodiode can be used to detect the timing of visual stimuli, which can be useful for calculating EMG activity changes relative to stimuli onsets. This is the last parameter inputted by the user.

Before the initiation of data measurement, a table (a flexible standard Matlab data type) with the variable name ‘trials’ is created in the Matlab workspace and pre-allocated with the number of rows corresponding to the number of sweeps to be recorded. Next, the function **EMGfigure** is called. This function sets up a figure where the data will be presented for on-line visualization. Separate panels distributed vertically on the screen are pre-allocated for each channel of data to be visualized (Figure 2).

**Figure 2.**
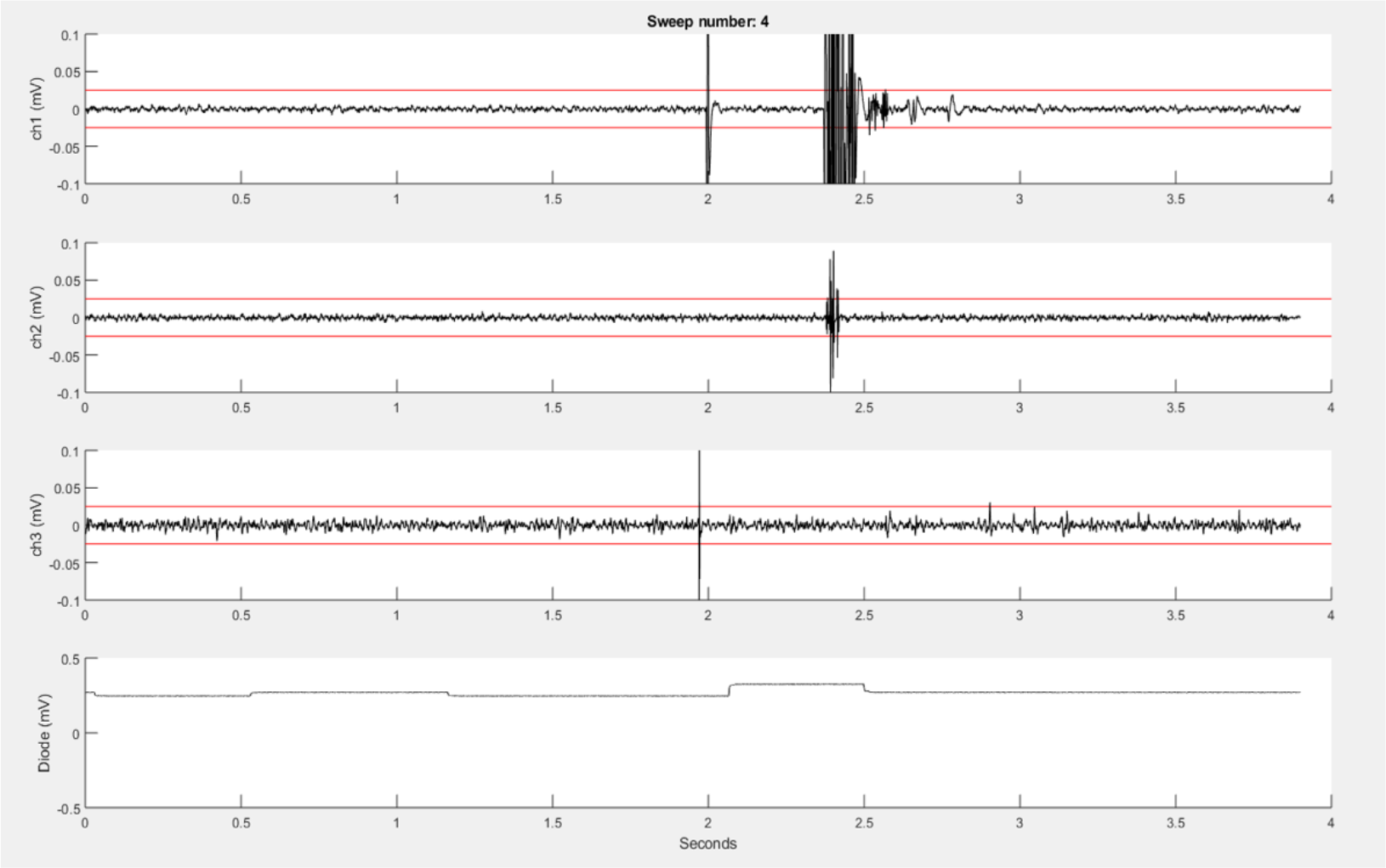
An example **recordEMG** live trace showing 3.9 s of a 4 second sweep recording three EMG channels and a photodiode channel. Channel 1 shows a MEP (at 2 s) and an EMG burst in response to a visual stimulus. Channel 2 shows a partial EMG burst associated with an incorrectly activated response. Channel 3 shows a TMS artefact. Channel 4 shows a photodiode signal with the increase at 2.1 s associated with the onset of the visual stimulus to which the participant responded. Red horizontal lines indicate .025 mV to serve as visual reference points for EMG and MEP amplitudes.

Using the ‘daq.createSession’ (Matlab Data Acquisition Toolbox) function, a data acquisition session object is created. The DAQ vendor, e.g. ‘ni’ for National Instruments, must be specified in the code, and appropriate drivers must be installed. Supported vendors can be found on the Mathworks website (https://www.mathworks.com/hardware-support/data-acquistion-software.html). The number of analog input channels that was inputted by the user is added to the session, and the sampling rate is set according to the specified parameters. EMG channels are added using the **addchannels** function. This function is written for a DAQ that can accommodate 9 channels (8 EMG channels and a supplementary input channel).

Additional channels can be added by modifying this function. Once the required number of recording channels are specified, an event listener is created using the Matlab built-in addlistener function. The listener retrieves and plots session object data to the figure using the callback function @**plotData**. The session begins operation with startBackground in order to allow the execution of additional code while the DAQ measurements are underway. The channel data are plotted in real-time during each sweep, and once a sweep is complete, the data are extracted from the plotted figure, using the **pulldata** function and stored in the ‘trials’ table according to each respective channel (column) and sweep number (row). Using this method, data are plotted with low latency, and then stored in the ‘trials’ table in the workspace until they are saved to the computer hard drive. The figure is cleared after each sweep for presentation of subsequent data. Each iteration passes sweep data to the ‘trials’ table in a successive row, and this process repeats for the set number of sweeps. This ordered procedure facilitates the timing of data visualization and data storage.

### 2) findEMG

This function automatically detects EMG burst, TMS, MEP, and CSP events.

#### Syntax

~~~
>>findEMG(‘filename’)
~~~

Specifying the filename as an input argument is optional.

#### Description

Default analysis parameters are specified below the header block similar to the **recordEMG** function. These include sampling rate (5000 Hz), EMG burst threshold (.3 V absolute threshold for detecting the presence of an EMG burst), EMG onset threshold (2 std of the mean EMG signal within a sweep is used to mark EMG onset/offset when an EMG burst is detected), TMS artefact threshold (.04 V) for the detection of TMS pulse artefacts in EMG traces, the time duration following the TMS artefact in which to detect an MEP (.1 s), the minimum time between the TMS artefact and MEP onset (.018 s), the pre-MEP time window (.1 s) used to calculate the root mean square of background EMG signal, and the threshold for the root mean square of EMG activity (.05 V) measured within this pre-MEP time window. Additional parameters are specified to facilitate the detection of EMG bursts within the same channels used to measure MEPs. These include the time prior to the TMS artefact (.005 s) and the estimated end of the MEP epoch in the EMG signal relative to the TMS artefact (.1 s). All these parameters can be tailored to the needs of an experimental analysis.

When calling **findEMG** the user either specifies an input filename or, if no input is provided, a dialogue box will open allowing the user to select an input file. The file must either have previously been saved as output from **recordEMG** or converted from another supported format. Functions for converting Acqknowledge^®^ (BIOPAC Systems Inc., Goleta, CA), EMGWorks^®^ (Delsys Inc., Natick, MA), and Signal^®^ (CED Ltd., Cambridge, England) data to the format needed to use **findEMG** are included with the toolbox. Additional conversion tools will be developed in the future, and this code will be made available through the GitHub repository.

Once the file is loaded, the number of channels in the data is automatically extracted from the trials table. The user is then prompted whether they would like the code to detect EMG bursts (1 for yes or 0 for no). If EMG bursts will be detected, the user is prompted to enter the number of the channels in which EMG bursts should be detected (e.g. [2] for channel 2, or [1 3] for channels 1 and 3). Separate prompts asks the user whether MEPs should be detected (1 for yes or 0 for no) and whether CSPs should be detected (1 for yes or 0 for no). If either is true, the user is prompted to enter the number of the channel that includes a recording of the TMS artefact without any muscle activity, e.g. from an EMG electrode placed over a cervical vertebra of the neck. If MEPs will be measured, the user is asked to enter the channel(s) in which MEPs should be detected (e.g. [1] for channel 1, or [2 3] for channels 2 and 3). If this value is set to 0, the code will search for TMS artefacts in the MEP channel. Similarly, if CSP will be measured, the user is prompted to enter the channel(s) numbers (e.g. [1] for channel 1, or [2 3] for channels 2 and 3). The entered MEP, CSP, and EMG burst channel(s) can overlap as all three metrics can be calculated within the same channel if desired.

If the user would like to analyze multiple files without being prompted to specify the inputs at the command line for each file, it is possible to forgo the command line prompts by defining the metrics and associated channel parameters within the code and setting the ‘use_command_line’ variable to 0. In this manner, filenames can quickly be passed to the **findEMG** code without further user input, which is convenient for batch file processing.

The main operations of the **findEMG** function include a series of automated routines for the detection of events in the data. First, the code evaluates whether a photodiode recording is present in the trials table. If so, the times of photodiode events are extracted using the **findDiode** helper function. This code can be modified to support the detection of events in other types of supplementary signals. Then, the **findEvent** helper function detects TMS artefacts, MEPs, EMG bursts, and CSPs in the appropriate channels specified by the user. In the case of TMS, TMS artefacts are located in the specified channel by searching for the artefact in the recorded EMG starting at the end of each sweep and scanning in reverse temporal order. This reverse approach ensures that MEP measurements are associated with the last TMS pulse in paired-pulse or multi-pulse TMS protocols as well as single-pulse protocols. MEP and CSPs are detected in time windows relative to the TMS artefacts using data from the user-specified MEP and CSP channel(s). All calculated metrics are outputted to the trials table in columns named with the appropriate channel number and metric listed in the **findEMG** header.

MEP detection depends on two user-defined parameters: 1) the duration of the epoch (.1 s) following the TMS artefact in which to search for a MEP, and 2) the minimal time elapsed following the TMS artefact at which a MEP onset can be marked (.018 s). Using the default parameters, MEP onsets are identified in the window between 18 and 100 ms following the TMS artefact. Matlab’s ‘findchangepts’ function is applied within this time window to identify points which minimize the sum residual squared error about the local mean. The default maximum number of change points detected is 10, and only the first and last points are used to define the MEP onset and offset. Once a MEP is identified, metrics are calculated including the MEP latency relative to the TMS artefact, maximum to minimum peak-to-peak amplitude, onset-to-offset duration, area, and root mean square of the EMG signal in the pre-MEP time epoch.

Additional automated routines in the **findEvent** helper function calculate the onset and offset times of EMG bursts. If EMG bursts will be detected in the same channel as MEPs, the EMG signal values from the time immediately prior to the TMS artefact through the subsequent MEP are assigned the mean of the signal for the entire sweep. In other words, any signal within this window are temporarily flattened to the mean. The values determining the beginning and end of this time window are specified in the parameters. This step makes it possible to ignore the TMS and MEP related changes in the EMG signal when searching for other patterns in the EMG activity. Once this step is complete, automated routines are performed to detect EMG bursts. These modified data are not stored and are only used for this procedure.

To detect EMG bursts, the code first detects whether the maximum rectified signal exceeds a fixed voltage threshold set in the parameters. If the code determines that a burst is present in the sweep, the function then combs the data for the first time point at which the voltage exceeds the user-defined EMG threshold, set as the specified number of standard deviations of the signal for the entire sweep (Greenhouse et al. 2015). This point is marked as the EMG burst onset for that channel for that sweep. To identify the burst offset, the script executes a similar procedure, but combing from the end of the trace in reverse temporal order. Once the EMG burst onset and offset are determined, the code also calculates the area of the rectified signal within the EMG burst epoch.

To detect CSPs, the code first determines whether MEPs were also detected. If so, the timepoint identified for MEP offset is also set as the CSP onset. The end point of the CSP, or CSP offset, is defined using a previously published method (King et al. 2006). In brief, Matlab’s findpeaks function is applied to the cumulative sum of the signal to identify the inflection point following CSP onset where the mean of the rectified signal starts to increase. The CSP offset is set to this time point. If the user wishes to calculate the CSP using the King et al. method without detecting MEPs, this can be done by only choosing to detect CSPs. However, the user should be aware that the calculated CSP onset time will differ depending on whether the MEP is detected or not detected in addition to the CSP.

Finally, the **findEMG** function outputs a file that contains the updated trials dataset with any desired TMS, MEP, RMS, CSP and EMG burst events within each sweep assigned to each appropriate channel. The ‘use_ui_save’ variable enables saving using the Matlab save dialogue box or, if disabled, will automatically save the output using the input filename appended with ‘_processed.’

### 3) visualizeEMG

This function displays a GUI for the visualization of EMG data for interactive review.

#### Syntax

~~~
>>visualizeEMG(‘filename’)
~~~

Specifying the filename as an input argument is optional.

#### Description

The **visualizeEMG** user interface was developed using GUIDE (Matlab’s GUI development environment) and enables data visualization to facilitate the identification of MEPs, EMG events, CSPs, TMS artefacts, and stimulus events. This function is associated with the visualizeEMG.fig file which determines the elements and graphical layout of the GUI.

When calling this function, the user loads a file that has been preprocessed with **findEMG**. The interactive components of the GUI are labeled in Figure 3. Advancing through sweeps in order is controlled through the toggle arrows at the bottom of the GUI. Alternatively, the user can jump to a specific sweep number by entering the desired sweep number in the text box. Zooming and panning capabilities are toggled on and off using the menu buttons at the upper left of the GUI and behave in the default manner.

**Figure 3.**
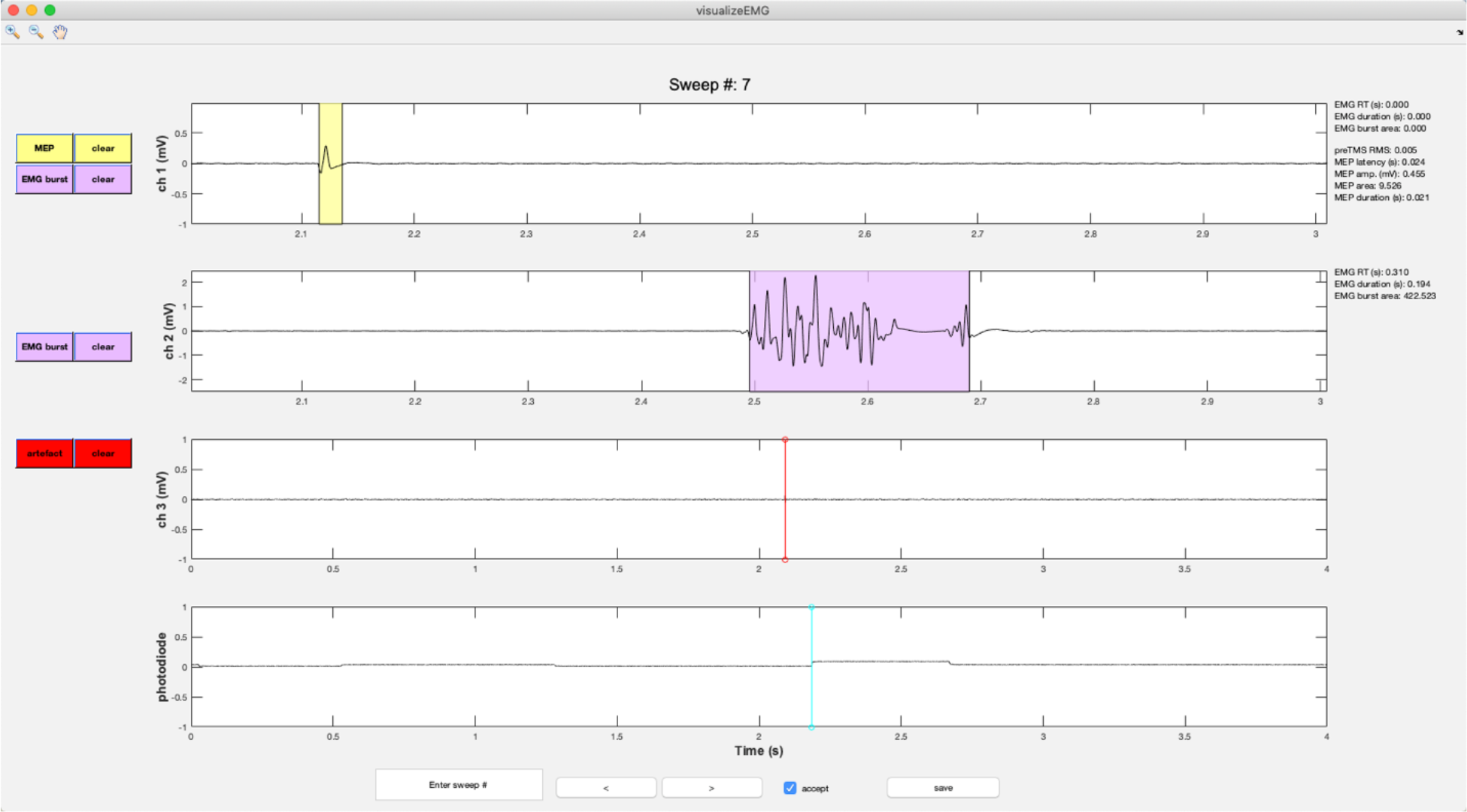
An example screenshot of the **visualizeEMG** GUI depicting three EMG channels and a photodiode channel. The sweep number is displayed at the top of the figure. Buttons on the left side allow the user to correct MEP indices and EMG bursts for each channel, as well as clear events that have been misidentified. The yellow shaded region indicates an MEP while the light magenta shaded region indicates an EMG burst. The red and cyan lines indicate TMS onset time and a photodiode event, respectively. If a sweep contains an EMG burst, EMG reaction time relative to the photodiode event, EMG burst duration, and EMG burst area are displayed in the right margin. If a sweep contains a MEP, preTMS root mean square, MEP onset latency relative to the TMS artefact, MEP amplitude, MEP area, and MEP duration are displayed. At the bottom of the figure, the user can jump to a specified sweep number, scroll between sweeps, accept or reject trials, and save the data. Data from channels 1 and 2 have been zoomed in to show trace detail using the zoom + tool in the upper left corner of the GUI.

Vertical lines are used to mark discrete events. These include TMS artefacts in red and stimulus onsets in magenta. EMG bursts are represented with light blue shaded regions, MEPs are represented with yellow shaded regions, and CSPs are represented with light green shaded regions (not shown).

Color-coded pushbuttons appear along the left side of the GUI, and their appearance depends on the number of channels recorded and types of events marked for display. A red ‘TMS’ pushbutton appears in the channel specified as containing TMS artefacts, and a red line indicates where an artefact has been detected by the **findEMG** code. Clicking the TMS pushbutton will activate crosshairs. Using the mouse to position the crosshairs over a new location in the TMS artefact channel and clicking will reset the TMS artefact position in the data. Only the position along the x-axis is used to manually redefine the location of the markers. The position along the y-axis is ignored. The GUI and ‘trials’ table immediately update to reflect any edits, and the ‘trials’ table automatically keeps a running tally of the total number of edits made to each sweep.

A similar editing procedure can be performed for MEP, EMG, and CSPs (Figure 4). However, these events are represented by colored patches that encompass the time from the event onset to the event offset. The buttons for each event type match the colors of the event patches. ‘MEP’ pushbuttons appear alongside MEP channels, ‘EMG’ pushbuttons appear alongside EMG bursts channels, and ‘CSP’ pushbuttons appear alongside CSP channels. Clicking any one of these buttons will activate the mouse-guided crosshairs that can be positioned over the location in the appropriate channel where a correction to the marked event is desired. Two mouse clicks are needed to define MEP, EMG, or CSP epochs. The first click indicates the onset, and the second click indicates the offset. Only the position of the cursor along the x-axis is used to define the manually entered onsets and offsets. The display and ‘trials’ table immediately update to reflect the corrected event markers, and these procedures can be repeated as many times as desired.

**Figure 4.**
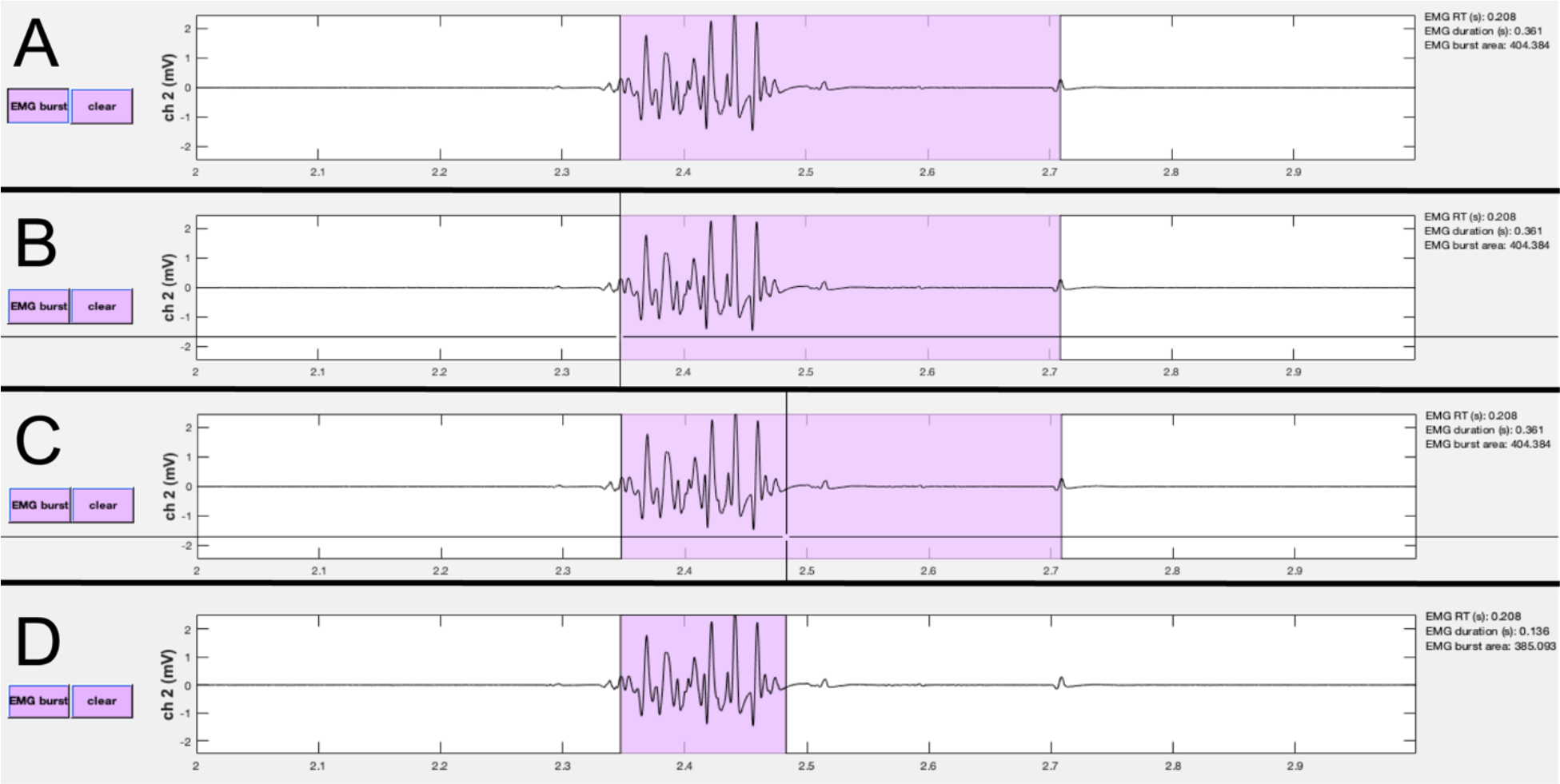
Events can be manually edited using the visualizeEMG buttons and mouse-guided cursor. In order to correct misidentified event onsets and/or offsets, the user presses the button on the left (A). By positioning the crosshair and clicking, the user selects the correct location of the burst onset (B) and offset (C). Lastly, the updated burst indices are recorded and the display is updated (D). Note, only the x-coordinates are recorded when choosing new indices. A similar method is executed to change MEP and CSP event indices.

Each event marker button is paired with a color-coded ‘clear’ button. Clicking any of these buttons will clear the event type from the actively displayed sweep. It is not necessary to clear an event before editing.

The user can also reject entire trials by unchecking the ‘accept’ checkbox at the bottom of the GUI. This is useful for excluding a sweep when data in all channels should not be included in subsequent analysis. A checked box indicates the trial will be flagged as acceptable, while an unchecked box indicates the trial is unacceptable for further analysis.

Metrics derived from the markers within each sweep are displayed in the right margin, adjacent to the plot displaying the data for each channel. These include the EMG burst onset time (EMG RT) relative to a stimulus such as a photodiode event or relative to the start of the sweep if no stimulus events are detected, the duration of the EMG burst, the area of the rectified EMG burst, root mean square of the EMG signal prior to TMS (RMS), MEP latency relative to the TMS artefact, MEP peak-to-peak amplitude, the area of the rectified MEP, the MEP duration, and the CSP duration. These values update for each respective channel and when any manual corrections are made to the data. The values are also stored in the trials table for subsequent analysis.

The ‘save’ button must be clicked before closing the GUI to save any changes made during the review process. Doing so will open Matlab’s save dialogue box. The default output file name is the input file name appended with ‘_visualized.’ Data can be saved and reopened at any point in the workflow as many times as needed. Further analysis of the data can be conducted using built-in Matlab functions (see GitHub repository for examples).

### Example Data & Analyses

Abbreviated example datasets (30 sweeps each) are included with the toolbox. These data were collected using Matlab 2017b on a Windows 10 Operating System, Delsys Bagnoli™ 8-channel EMG System interfaced with a National Instruments PCI-6220 card, and a Magstim® 200-2 single pulse TMS stimulator. EMG signals were amplified x1000, bandpass filtered online between 20 and 450 Hz, and sampled at 5000 Hz. Adapted versions of **recordEMG** were used to visualize and store the data during all measurements.

Data are included from four experiments: 1) a TMS short intracortical inhibition (SICI) protocol with conditioning pulses at 50%, 65%, 80%, 95%, and 110% active motor threshold and test pulses at 115% resting motor threshold, 2) a CSP protocol with tonic contraction maintained at 25% MVC and TMS administered at 115% resting motor threshold, 3) a unimanual Stop Signal Task with maintained tonic contraction of a task-irrelevant muscle (no TMS was administered during the acquisition of these data), 4) a Delayed Response Task which included TMS pulses administered at 115% resting motor threshold delivered at one of two time points on a subset of trials: either an inter-trial baseline or during a preparatory delay period 100 ms prior to an imperative Go stimulus. These datasets were collected at different recording sessions using EMG from the first dorsal interosseous (FDI) muscles of the hands.

To assess the robustness of our analysis tools, each example dataset was analyzed using the **findEMG** automated event detection with the appropriate channels indicated for EMG burst, TMS artefact, MEP, and CSP detection. For the SICI experiment, TMS artefacts were detected in channel 2, and MEPs were detected in channel 1. For the CSP experiment, TMS artefacts were detected in channel 3 and both MEPs and CSPs were detected in channel 1, with resting EMG in channel 2. For the stop signal task, only EMG burst events were detected in channel 1. For the delayed response task, EMG burst events were detected in channels 1 and 2, TMS artefacts were detected in channel 3, and MEPs were detected in channel 1. Otherwise, parameters were matched across datasets, and the default settings were used.

Following the automated detection of events with **findEMG**, the **visualizeEMG** code was used to manually inspect the data and make corrections where desired. The EMG event numbers and statistics were calculated after running **findEMG** alone and again after visual inspection with **visualizeEMG**. The values are presented in Table 1.

**Table 1.**
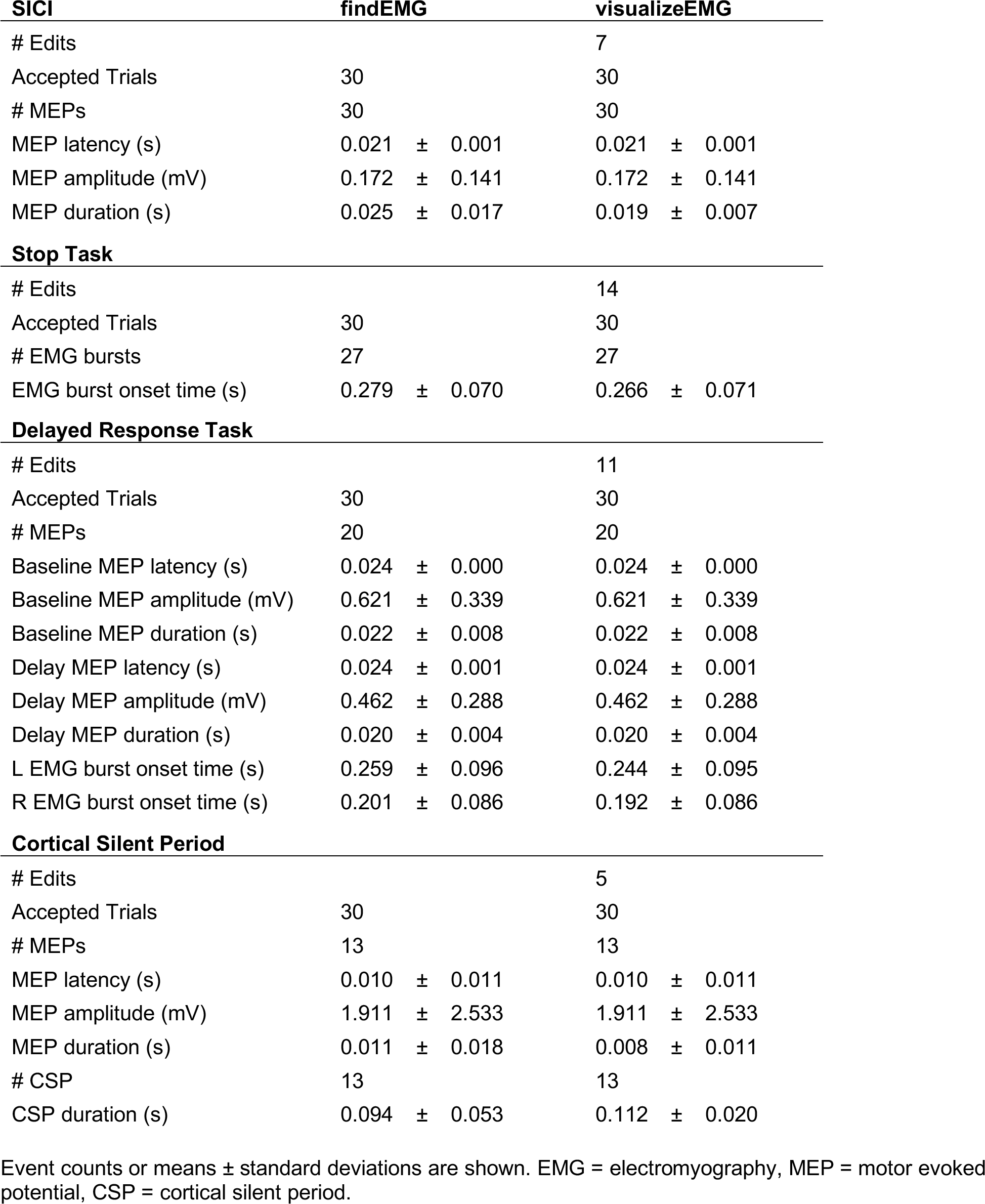
Event statistics for automated and interactive event detection.

Edits were made to all types of the example data suggesting that interactive visualization is useful for refining the automated event detection **findEMG** algorithms regardless of the type of data being analyzed. Notably, certain features of the data were more likely to be edited during visualization than others. For example, the overall number of detected events did not differ between the automated and interactive visualization of the data. In contrast, the EMG burst onset times determined from the automated detection algorithm tended to be later than those identified with interactive visual inspection of the data. The adjustments to EMG burst onsets were made on a subset of trials as indicated by the fact that edits were only made on a subset of the sweeps. The estimated CSP duration was longer following visualization than with the automated routines alone. This was because the automated detection of the CSP offset failed and assigned a time point before the end of the CSP on five sweeps, when the amplitude of the signal associated with the tonic contraction decreased. Interestingly, MEP latencies and amplitudes did not differ between the automated and interactive procedures although MEP durations were shorter following visualization of the data indicating the MEP offset was occasionally overestimated.

## Discussion

The VETA Matlab toolbox is a versatile suite of tools for the recording and analysis of EMG data with an emphasis on MEP detection and measurement. The toolbox simplifies essential steps of the EMG data collection and analysis pipeline. Low-latency recording and on-line visualization of EMG data is supported with the **recordEMG** tool. Automated detection and calculation of common metrics are supported with the **findEMG** tool. The off-line visualization and editing of events are supported with the **visualizeEMG** tool. These tools capitalize on built-in features of Matlab and are designed to be accessible to beginner Matlab users.

Example datasets accompany the toolbox and include SICI measurements acquired outside a behavioral task, CSP measurements also acquired outside a behavioral task, EMG measurements taken during the performance of a Stop Signal Task without TMS, and EMG measurements taken during the performance of a Delayed Response Task with MEPs elicited during specific task intervals. These datasets were used to evaluate the functionality of the automated detection and visualization tools and demonstrate their basic usage, recommended workflow, and successful application. Our findings from the processing of these data indicate that the event detection algorithms are robust across a variety of experiments including automated detection of EMG bursts occurring in close temporal proximity to MEP measurements within the same EMG recording channel. Our findings further indicate that MEP duration, CSP duration, and EMG burst onset calculations benefit from interactive visualization and manual editing with the **visualizeEMG** tool.

Several features of VETA differentiate this toolbox from others. While the **recordEMG** tool stands alone, the underlying code is easily adapted for use in combination with Psychtoolbox (Brainard et al., 1997) to measure EMG and MEPs during the simultaneous presentation of behavioral stimuli and response collection. A major advantage of combining these tools is the capability to collect EMG data and control experimental stimuli from a single computer, alleviating challenges often met when interfacing multiple computers. The benefits of a single stimulus presentation and data recording platform may be particularly valuable when conducting experiments in challenging settings such as the clinic or operating room. Example code for the administration of a Delayed Response Task that includes stimulus presentation, EMG recording, and the delivery of timed TMS pulses is included with the toolbox. This code is commented to facilitate editing for other task paradigms and a version of this code was used to create the Delayed Response Task example dataset that accompanies the toolbox. Moreover, while the **recordEMG** tool was developed with EMG data collection in mind, this tool will work for any type of data that can be acquired using Matlab’s Data Acquisition Toolbox and is by no means restricted to EMG.

VETA capitalizes on the flexibility of Matlab’s table data type to offer a standard format for storing EMG data. The included example datasets illustrate the recommended use of the table for storing EMG data. Moreover, each file generated by **recordEMG** includes an additional ‘subject’ variable for storing subject characteristics, and simply removing this variable anonymizes the data for sharing. This approach offers a straightforward solution to storing useful subject information alongside the data and supports quick data anonymization when desired.

The VETA toolbox is limited in its current form. One limitation is that only certain DAQ vendors are supported through Matlab’s Data Acquisition Toolbox. While this prohibits the use of **recordEMG** with unsupported vendors, it is still possible to apply the **findEMG** and **visualizeEMG** tools for the analysis of data collected using other software. Data conversion tools accompany the toolbox and support conversion of common EMG data formats to the table Matlab data type used by the VETA toolbox.

These currently include Acqknowledge^®^ (BIOPAC Systems Inc., Goleta, CA), EMGWorks^®^ (Delsys Inc., Natick, MA), and Signal^®^ (CED Ltd., Cambridge, England) formats, and additional formats will be supported in the future. Following conversion of these data formats to the VETA ‘trials’ table, **findEMG** and **visualizeEMG** will work with no additional required changes to the data. Finally, although the toolbox was developed for EMG and TMS, the tools lend themselves to the analysis of other types of biosignal processing, and other types of data may be supported in the future.

All code and example data are freely available through the GitHub repository (https://github.com/greenhouselab/VETA).

## Conclusion

The VETA toolbox simplifies the EMG data collection and analysis pipeline. Three main tools are included. The first performs online visualization and recording of EMG data. This tool integrates well with existing Matlab stimulus presentation toolboxes to facilitate the combination of EMG, TMS, and behavioral testing. The second tool automatically detects a variety of signal events including TMS artifacts, EMG bursts, MEPs, and CSPs, and calculates commonly reported metrics. The third tool is an interactive GUI for reviewing and editing data. Data are stored in a format to expedite further analysis and promote open science practices.

## Acknowledgements

I Greenhouse is supported by NIH National Center for Advancing Translational Science TR002370-01. We thank Mike Claffey for developing a precursor toolbox and Florent Lebon and Jan Wessel for sharing sample data to build initial conversion tools.

